# A toolbox for studying cortical physiology in primates

**DOI:** 10.1101/2021.08.04.455066

**Authors:** Karam Khateeb, Julien Bloch, Jasmine Zhou, Mona Rahimi, Devon J. Griggs, Viktor N. Kharazia, Min N. Le, Ruikang Wang, Azadeh Yazdan-Shahmorad

## Abstract

Lesioning and neurophysiological studies have facilitated the elucidation of cortical functions and mechanisms of functional recovery following injury. Clinical translation of such studies is contingent on their employment in non-human primates (NHPs), yet tools for monitoring and modulating cortical physiology are incompatible with conventional NHP lesioning techniques. To address these challenges, we developed a toolbox demonstrated in seven macaques. We introduce the photothrombotic method to induce focal cortical lesions alongside a quantitative model for the design of lesion profiles based on experimental needs. Large-scale (∼5 cm^2^) vascular and neural dynamics can be monitored and lesion induction can be validated *in vivo* with optical coherence tomography angiography and our electrocorticographic array, the latter of which also enables testing stimulation-based interventions. By combining optical and electrophysiological techniques in NHPs, we can enhance our understanding of cortical functions, investigate functional recovery mechanisms, integrate physiological and behavioral findings, and develop treatments for neurological disorders.

## Introduction

The primate neocortex is responsible for a variety of complex tasks and behaviors, including long-term memory storage, sensory processing, and movement. Historically, both lesioning and neurophysiological studies have been critical for elucidating functions of specific cortical regions (Ferrier, 1876) such as somatosensory (Borich et al., 2015; Brinkman et al., 1985; Gerlai et al., 2000), visual (Humphrey, 1974; Wurtz and Goldberg, 1972), auditory (Heffner and Heffner, 1986), and posterior parietal (Murphy et al., 2016; Vallar et al., 1994) cortices. Recently, such strategies have also been employed to investigate mechanisms of plasticity and recovery following injury (Friel et al., 2007; Harrison et al., 2013; Kaeser et al., 2010; Liu and Rouiller, 1999; Nudo and Milliken, 1996; Nudo et al., 1996; Padberg et al., 2010; Pons et al., 1988; Xerri et al., 1998a). The study of these phenomena in non-human primate (NHP) models with strong evolutional relevance to the human cortex is critical for understanding fundamentals of cortical physiology and designing novel clinical treatments for cortical injury.

Typical strategies for investigating *in vivo* NHP cortical physiology include either monitoring or perturbing the native cortical activity then correlating neural activity with behavior. Neural activity is frequently monitored through electrical recording, or calcium imaging, while perturbations include lesioning, electrical stimulation, or optogenetic manipulation (Tremblay et al., 2020). Previously, we developed a large-scale interface enabling optogenetic neuromodulation in concert with simultaneous electrical recording (Ledochowitsch et al., 2015a; Yazdan-Shahmorad et al., 2015, 2016, 2018a, 2018b). However, to the best of our knowledge, there is no single unifying paradigm through which the full spectrum of strategies can be combined for the unhindered investigation of cortical physiology in NHPs. Conventional lesioning techniques lack compatibility with tools for monitoring and modulating cortical physiology and lack flexibility in controlling lesion location and extent. Here, we integrate a versatile focal ischemic lesioning technique with large-scale monitoring of cortical vascular dynamics, and electrophysiological recording and stimulation. Importantly, with large-scale stable optical access, this toolbox can be combined with optical stimulation and imaging techniques such as optogenetics and calcium imaging, respectively.

Commonly utilized NHP cortical lesioning techniques are challenging to employ. One method of lesioning cortex in NHPs is through middle cerebral artery occlusion (Maeda et al., 2005; Virley et al., 2004). Because middle cerebral artery occlusion is used to mimic ischemic stroke as observed in the clinic, the resulting lesions are broad and are constrained only to regions downstream of the middle cerebral artery. Moreover, obtaining access to the middle cerebral artery for occlusion requires complex surgical intervention regardless of the occlusion technique. Similarly, common focal cortical lesioning techniques such as endothelin-1 (Dai et al., 2017; Teo and Bourne, 2014), electrocoagulation (Nudo et al., 2003; Xerri et al., 1998b), ibotenic acid (Kaeser et al., 2010; Liu and Rouiller, 1999), cooling (Brinkman et al., 1985), and aspiration (Heffner and Heffner, 1986; Padberg et al., 2010; Pons et al., 1988) also involve technically challenging surgical procedures that are susceptible to variability across animals. Importantly, these techniques la ck compatibility with tools for monitoring cortical physiological dynamics during lesion formation and recovery.

When conducting lesion studies in NHPs, it is critical that lesions are validated *in vivo* for studies involving long-term post-lesion behavioral and physiological assessment. Current lesion validation methods primarily consist of behavioral assessment and post-mortem histological analysis (Kaeser et al., 2010; Liu and Rouiller, 1999; Murata et al., 2008). Methods of *in vivo* lesion validation, in addition to behavioral assessment, include magnetic resonance imaging (MRI) (Le Friec et al., 2020), or visual assessment of tissue blanching due to devascularization (Nudo and Milliken, 1996; Nudo et al., 1996; Xerri et al., 1998a). While MRI validation can provide both anatomical and physiological assessment of lesion induction, it exhibits poor spatial and temporal resolution (Baran and Wang, 2016). Moreover, visual assessment can result in unreliable estimation of lesion boundaries (Xerri et al., 1998a). The reliance on behavioral and post-mortem histological analysis alone is deficient of direct anatomical and physiological lesion validation during the lifetime of the animal. Ideally, lesion validation would be accomplished reliably, and in a manner that allows for monitoring of anatomical and physiological changes over the course of recovery.

In addition to these techniques, the investigation of cortical neurophysiological dynamics can play an important role in enhancing our understanding of cortical physiology and in developing neurorehabilitative strategies. The monitoring of neurophysiological dynamics in neuroscience is often accomplished through electrical stimulation and recording of neural activity. For example, intracortical microstimulation of primary motor cortex following a lesion has been crucial in elucidating the remapping of cortical somatotopic representations (Nudo and Milliken, 1996; Nudo et al., 1996). In other cortical regions, electrophysiological recordings have also been used to monitor neural activity following lesion induction (Padberg et al., 2010; Schmid et al., 2009). The incompatibility of the lesioning methods used in these studies with recording and stimulation techniques, however, prevents the monitoring of neural activity simultaneously during lesion formation. Ideally, neural activity would be recorded before, during, and after lesion formation, while stimulation-based interventions could be tested after lesioning.

In this study, we addressed the technical shortcomings of studying cortical physiology in NHPs. We successfully demonstrate the combination of a photochemical lesion induction technique with a controlled spatial profile, an *in vivo* lesion validation method with high spatial resolution, and the ability to simultaneously monitor the underlying neural activity and blood flow as lesions form at a large scale (∼5 cm^2^). This toolbox also offers the capability to test stimulation-based interventions. Moreover, the sizes of focal lesions induced with our toolbox are comparable to previously reported focal lesions capable of eliciting behavioral deficits (Murata et al., 2008; Padberg et al., 2010). Importantly, the tools presented here are compatible with previously established interfaces offering large-scale access for optical and electrical stimulation as well as imaging (Griggs et al., 2021b; Macknik et al., 2019; Yazdan-Shahmorad et al., 2016), providing unparalleled access to the primate cortex. Through this integrative approach, the large-scale monitoring of cortical network activity and reorganization concurrently with vascular dynamics expands the questions and interventions we can explore, profoundly impacting the development of neurorehabilitative therapies.

## Results

To address the challenges of studying cortical physiology in NHPs, we developed a toolbox that allows for targeted focal lesioning, large-scale imaging of cortical blood flow, and large-scale electrophysiological recording and stimulation. Our toolbox is comprised of the photothrombotic technique for inducing focal ischemic lesions, optical coherence tomography angiography (OCTA) imaging for *in vivo* lesion validation, histological validation, a computational model for designing lesion profiles, and electrocorticographic (ECoG) recording during lesion formation. We implemented the photothrombotic technique (Figure 1) in 7 adult macaques (monkeys A-G) and validated the disruption of cortical blood flow *in vivo* using OCTA imaging as described in the STAR Methods. Lesions were induced by illuminating through an opaque apertured mask (Figure 1A) in a 25 mm diameter cranial window with an uncollimated white light source following intravenous infusion of the photoactive dye Rose Bengal (Figure 1B). Photoactivation of Rose Bengal in the cerebral vasculature results in the release of reactive oxygen species, subsequent endothelial cell damage, platelet activation, and vascular occlusion due to the formation of thrombi. We tested the effects of a range of aperture diameters (0.5 to 2 mm) and illumination intensities (0.1 to 18.6 mW) on lesion size. To test these variables, we used an opaque mask with multiple apertures of various diameters dispersed throughout the cranial window (Supplementary Figure 1). A single uncollimated white light source was used to illuminate through the mask apertures such that the light intensity was highest through centrally located apertures. Through this approach we were able to test a variety of illumination parameters while reducing the number of animal subjects, time under anesthesia, and the number of light sources to practical levels.

**Figure 1.**
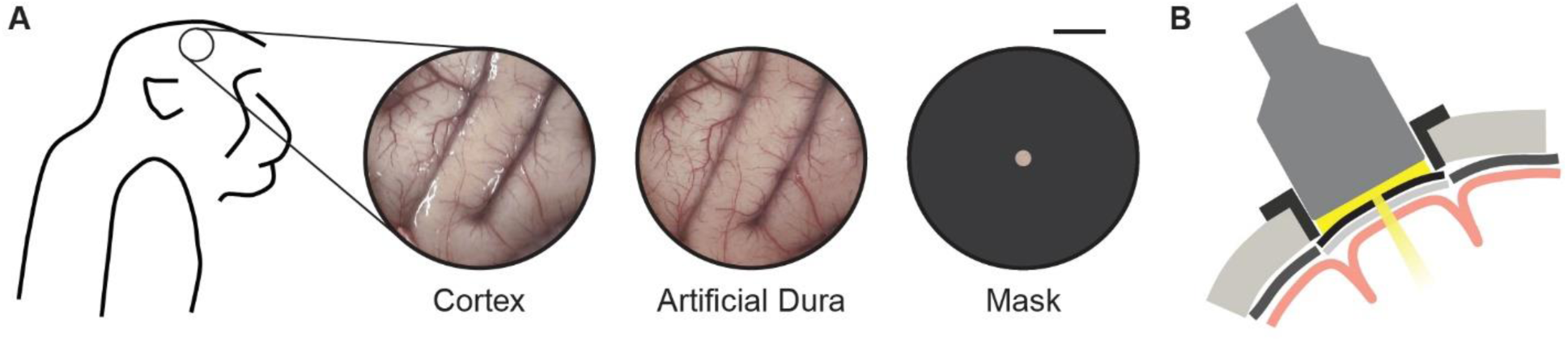
Schematic of photothrombotic technique application to induce focal cortical ischemic lesions. (A) Following a 25 mm diameter circular craniotomy, a thin, transparent artificial dura was placed over the exposed cortical surface. Next, an apertured mask was placed on top of the artificial dura. Scale bar is 5 mm. (B) Coronal schematic of light illumination through the apertured mask following intravenous Rose Bengal infusion. Portions of this figure have been adapted for inclusion in this manuscript from Khateeb et al., 2019 with permission.

### Validation of Neuronal Cell Loss by Histology and Immunohistochemistry

Approximately 4 hours after the end of illumination, animals were perfused with 4% paraformaldehyde (PFA) and the brains were extracted. Prior to histological staining of coronal slices, we observed bright pink areas corresponding with the illuminated regions (Figure 2A), suggesting that the formation of thrombi occluded the vasculature, entrapping the pink-colored Rose Bengal even after perfusion. We then stained coronal sections of the brains for Nissl bodies (Figure 2B,C). In illuminated regions, we observed areas characterized by well-defined borders that lacked Nissl staining, demonstrating cell loss in those regions (Figure 2B,C). We identified these areas as lesions. To test for neuronal cell loss, we performed NeuN staining and observed similarly defined borders in the illuminated regions (Supplementary Figure 2). Depending on aperture sizes and illumination intensities, lesion depths extended through all cortical layers. To gain perspective of the relative locations and sizes of the lesions, we reconstructed the lesions in three-dimensional space using the coronal Nissl-stained sections (Figure 2D-H). With this method, we measured lesion volumes ranging between 1.7 mm^3^ and 35.4 mm^3^. We observed 22 histologically detected lesions out of 30 illuminated areas across monkeys B-E. Notably, high intensity illumination through centrally located apertures more consistently resulted in histologically detected lesions (lesion volumes ranging from 10.1 mm^3^ to 31.2 mm^3^, median of 19.6 mm^3^) compared to lower intensity illumination in peripheral apertures (volumes 0 mm^3^ to 35.4 mm^3^, median of 2.9 mm^3^).

**Figure 2.**
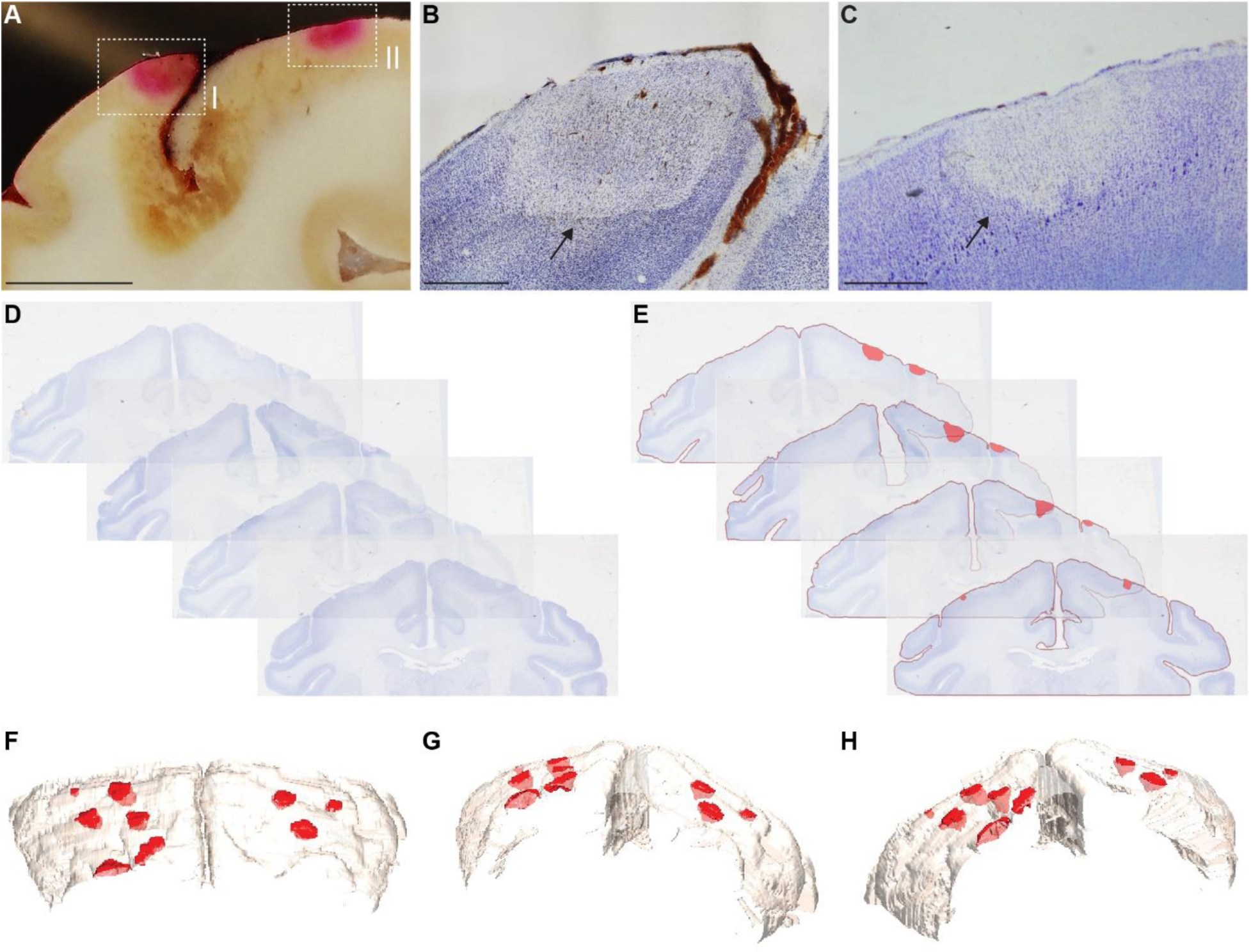
Histological lesion validation and reconstruction. (A) Unstained coronal section from monkey A. Pink regions corresponding with illuminated regions highlighted in boxes I and II indicate the presence of Rose Bengal entrapped in the cortical microvasculature. Scale bar is 5 mm. (B,C) Coronal Nissl-stained slice adjacent to the slice shown in panel (A), showing cell loss in the region encapsulated in box I (B) and II (C) and indicated by the black arrows. Scale bars are 1 mm. (D,E) Reconstruction of induced lesions in 3D space was done by first co-registering coronal Nissl-stained slices (D), followed by identification of lesion boundaries (E). (F-H), 3D reconstruction of lesions in monkey C from three different angles. Portions of this figure have been adapted for inclusion in this manuscript from Khateeb et al., 2019 with permission.

### Optical Coherence Tomography Angiography for Large-Scale Blood Flow Imaging and *In Vivo* Lesion Validation

The extent of optical access afforded by our toolbox enables for the application of large-scale imaging techniques such as OCTA imaging. OCTA is a non-invasive angiographic technique that is capable of imaging functional vascular networks within tissue beds *in vivo*. OCTA has been demonstrated to detect inflammatory conditions in skin (Deegan and Wang, 2019; Deegan et al., 2018a), eyes (Kashani et al., 2017), and brain (Li et al., 2018; Park et al., 2018). While post-mortem histological analysis is the predominant method of measuring focal lesion size, we introduce OCTA as an *in vivo* tool to image blood flow in the cortical microvasculature, validate lesion induction, and measure ischemic lesion sizes. We imaged several 9 mm x 9 mm cortical regions subjected to photothrombotic ischemia.

OCTA imaging 3 hours after illumination revealed clear localized disruptions in cortical blood flow compared to baseline images in the illuminated regions (Figure 3, Supplementary Figure 3), as evidenced by a lack of functional blood vessels in regions interrogated by the photothrombotic illumination. Thus, we demonstrated the ability to validate lesion induction *in vivo* through OCTA imaging.

**Figure 3.**
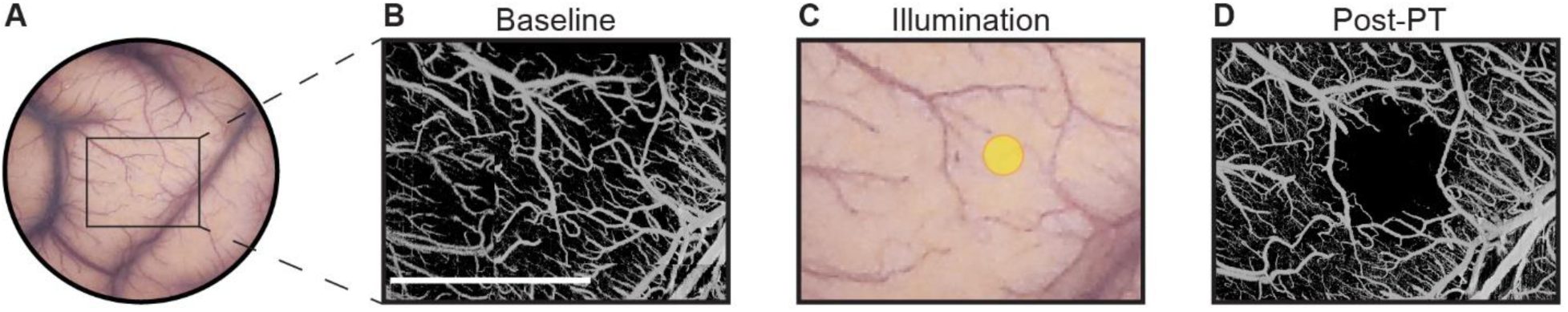
Optical coherence tomography angiography (OCTA) validation of lesion induction. (A) Surface of sensorimotor cortex through an artificial dura in monkey C. (B) OCTA imaging of the rectangular area prior to lesion induction. Scale bar is 5 mm. (C) Illumination of a cortical region of interest indicated by the yellow circle following intravenous Rose Bengal infusion. (D) OCTA imaging 3 hours post-photothrombosis. Portions of this figure have been adapted for inclusion in this manuscript from Khateeb et al., 2019 with permission.

We then compared OCTA and histological lesion validation modalities (Figure 4A-C). Not all illuminated regions resulted in lesions detected through both modalities. Across monkeys B through E, we illuminated 30 regions to induce photothrombosis. 26 of the illuminated regions were imaged within the OCTA field of view (FOV), of which 16 lesions were detected through OCTA imaging. 15 of the 16 OCTA-detected lesions were later confirmed histologically with one lesion undetected histologically. Similar to histology, illumination through centrally located apertures corresponding with higher light intensities more consistently resulted in lesions detected with OCTA compared to peripheral apertures with lower light intensities. Additionally, 8 of the 10 imaged regions in which no lesions were detected in OCTA images corresponded with regions occupied by large blood vessels. Of those 8 regions, only 3 resulted in lesions detected through histology (3.7 to 17.6 mm^3^). Of the 22 histologically detected lesions, 15 were detected through OCTA. Histologically detected lesions within the OCTA FOV that were not detected in OCTA images (4 lesions) were likely a result of illumination of large vessels (3 lesions), or the lesion may have been too small to be detected via OCTA imaging (1 lesion, 2.83 mm^3^). Notably, OCTA detection of lesions was unsuccessful in all instances in which the illuminated region coincided with large blood vessels, regardless of histological detection (Supplementary Figure 4). Because of the inconsistencies of large vessel illumination due to differences in optical and biological properties, we excluded data corresponding with large vessel illumination from further analysis.

**Figure 4.**
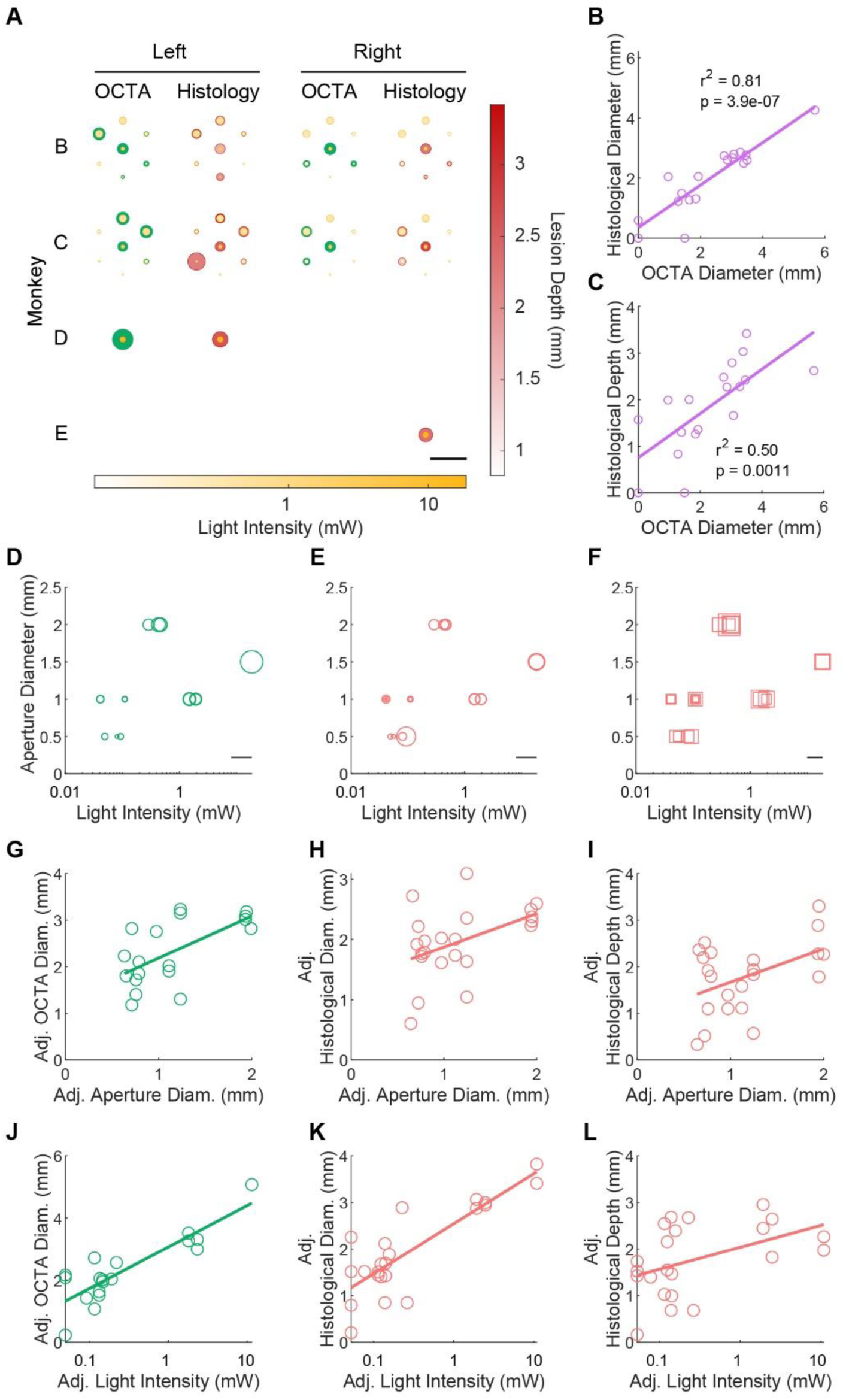
Comparison of OCTA and histological lesion validation and effect of illumination parameters. (A) Schematic demonstrating the OCTA-measured lesion diameters (green circles) in the illuminated regions (yellow circles) of both hemispheres for monkeys B-E. The shading of the yellow circles indicates illumination intensity. The histologically measured lesion diameters are also shown (pink circles), with the shading indicating lesion depth. (B) OCTA-measured and histologically measured lesion diameters were highly correlated (0.81 r-squared, p = 3.94e-7). (C) OCTA-measured lesion diameters also correlated with histologically measured lesion depth (0.50 r-squared, p = 0.0011). (D) Combined effect of aperture diameter and light intensity on OCTA-measured lesion diameter. Circular marker diameters represent OCTA-measured lesion diameter. Scale bar is 5 mm. (E) Combined effect of aperture diameter and light intensity on histologically measured lesion diameter, where the diameter of the circular markers represents histologically measured lesion diameter. Scale bar is 5 mm. (F) Combined effect of aperture diameter and illumination intensity on histologically measured lesion depth. The width of the square markers denotes histologically measured lesion depth. Scale bar is 2 mm. (G-L), Partial regression plots demonstrating the effects of aperture diameter and light intensity via a multivariate linear model on OCTA-measured lesion diameters (0.85 r-squared, p = 6.5e-7) (G,J), histologically measured diameters (0.81 r-squared, p = 1.8e-7) (H,K), and histologically measured depths (0.49 r-squared, p = 0.0017) (I,L). In panels G-L, the partial regressions are shown with the adjusted variables.

We then quantitatively assessed OCTA-measured lesion diameters as *in vivo* proxy measurements of histologically measured lesion diameters (Figure 4B). We observed a positive linear relationship between OCTA-measured lesion diameters and histologically measured lesion diameters, with an r-squared value of 0.81 (p = 3.9e-7; Figure 4B). Because OCTA imaging provides depth information only down to about 600 µm, we assessed the correlation between histologically measured lesion depths and OCTA-measured lesion diameters (Figure 4C). We observed a positively correlated linear relationship between histological lesion depth and OCTA-measured lesion diameter with an r-squared value of 0.50 (p = 0.0011; Figure 4C). Interestingly, we also observed a significant positive correlation between both histologically measured lesion depths and diameters (0.65 r-squared; p = 5.3e-6; Supplementary Figure 4). Importantly, our results demonstrate that OCTA-measured lesion diameter is a strong predictor of histologically measured diameters.

### Effect of Illumination Parameters on Lesion Size

We examined the combined contributions of both aperture diameter and illumination intensity as predictors of lesion size through multivariate linear regression analysis (Figure 4G-L, Supplementary Table 1). As the Beer-Lambert law (Bouguer, 1729) predicts an optical path length with logarithmic relation to light intensity, we used the log-transformed light intensity as a predictor. We found that a model accounting for both sets of illumination parameters accounts for a large portion of the variance in OCTA-measured diameters with an r-squared value of 0.85 (p = 6.5e-7; Figure 4D,G,J). In predicting histologically measured lesion diameters, the combined illumination parameters yielded an r-squared value of 0.81 (p = 1.8e-7; Figure 4E,H,K). For the prediction of histologically measured lesion depths based on both aperture diameter and light intensity, we observed an r-squared value of 0.49 (p = 0.0017; Figure 4F,J,L). Therefore, both illumination parameters serve as significant determinants of lesion size, and the consideration of their combined effects is critical for predicting lesion sizes.

### Prediction of Lesion Size by Simulation of Light Propagation through Cortical Tissue

To extend the predictability of lesion sizes based on illumination parameters beyond those tested in this study, and to be able to predict both lesion diameter and depth with a single biophysically inspired model, we developed a simulation-based computational model. Photothrombotic lesions arise due to thrombi formation and subsequent cell death mediated by the topography of the underlying microvasculature. As such, we developed a two-stage modeling process in which we simulated photons penetrating brain tissue (Figure 5S) to generate a profile of the spatial fluence distribution (Figure 5B) and transformed fluence contours to recreate the lesion shapes observed through histology (Figure 5C,D).

**Figure 5.**
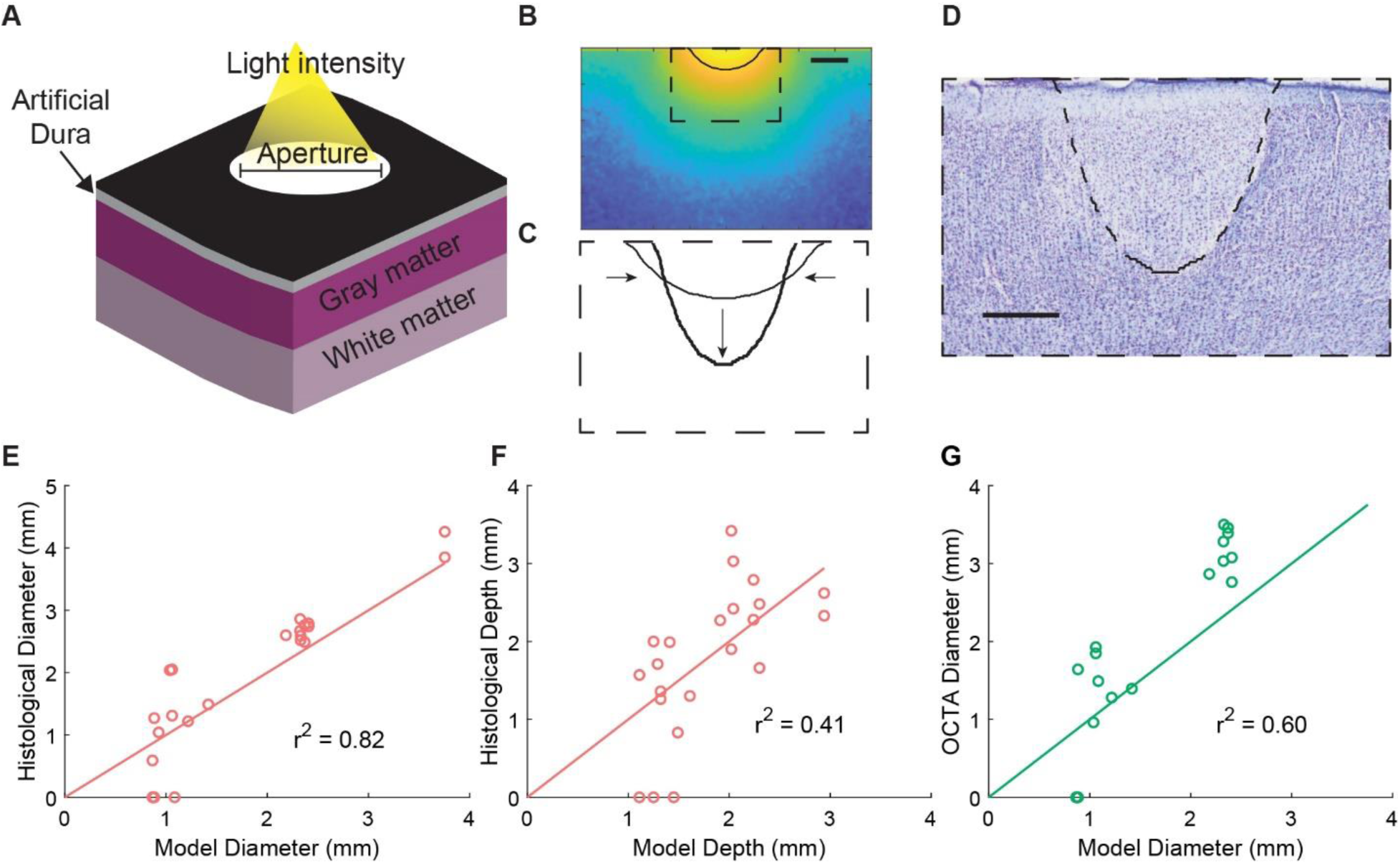
Prediction of lesion size by simulation of light propagation through cortical tissue. (A) Schematic of simulated cortical volume with our experimental setup. An uncollimated light beam passes through an aperture and a transparent artificial dura (0.5 mm thick), into gray and white matter of a virtual cortical medium. Gray matter thickness is 2.5 mm. (B) A contour is identified from the light profile matching the light intensity threshold (19.9 µW/mm^2^) most closely matching the lesions. Scale bar is 100 µm. (C) The light intensity contour is scaled to generate a biological lesion contour. Scaling factors were obtained through regression on our dataset of simulated lesion dimensions and corresponding histologically measured lesions. (D) Predicted lesion contour overlayed on a coronal Nissl-stained slice of a corresponding lesion from monkey B. Scale bar is 50 µm. (E-G), Modeled lesions accurately predict histologically measured lesion diameters (0.82 r-squared) (E) depths (0.41 r-squared) (F), and OCTA-measured diameters (0.60 r-squared) (G).

To model the illuminated cortical tissue, we constructed a virtual optical illumination and medium model of our experiments, consisting of a light source and mask, artificial dura, gray matter, and white matter (Figure 5A). The entire volume spanned a size of 8 mm x 8 mm x 5.6 mm (width x width x height) and was constructed in the Monte Carlo eXtreme (MCX) software (Fang and Boas, 2009). In this volume, uncollimated light exiting our compound light source (see STAR Methods) passes through the artificial dura, gray matter, and white matter. By simulating stochastic photon transmission through this medium based on different illumination parameters, we can obtain spatial fluence distributions over the volume. As the photothrombotic lesioning method relies on the interaction between light and Rose Bengal to occlude blood flow, this spatial distribution can describe the effect of illumination parameters on lesion induction.

The optical properties of gray and white matter scattering and absorption coefficients had wide ranging values previously reported in literature (Gottschalk, 1992; Yaroslavsky et al., 2002). Therefore, we ran a grid-search of Monte Carlo simulations of 1 million photons propagating through a virtual medium with these optical properties within the reported ranges to find a best fit to our data. For each set of optical properties in the grid we ran a simulation for our experimental aperture diameter and illumination intensity combinations. A spatial fluence distribution threshold was then identified (Figure 5B) for each set which maximized the overlap (see Equation 2, Methods) between the simulated lesions and the corresponding lesions obtained through histology. Using this method, we identified the best optical properties as a gray matter absorption coefficient of 0.395 mm^-1^, gray matter scattering coefficient of 53.6 mm^-1^, white matter absorption coefficient of 0.09 mm^-1^, white matter scattering coefficient of 54.066 mm^-1^, and a spatial fluence distribution threshold for lesion induction of 19.9 µW/mm^2^.

From the identified best-matching light simulations we quantified the degree to which they predict lesion shapes and sizes. We observed that while the simulation results qualitatively align with the lesion profiles, the light simulation alone was not adequate for explaining the observed lesions. By using the maximal depth and average diameter of individual fluence threshold contours of the simulation as predictions of their corresponding lesion profiles, we observed that the contours resulted in an r-squared value of -0.04 for lesion depth and 0.32 for diameter. These results indicate that the extent of induced photothrombotic ischemic lesions *in vivo* is not simply a function of the fluence profile in the cortex but is likely also governed by biological factors such as vascular topography.

We therefore incorporated a scaling process to transform the light intensity contours to the biological lesion profiles observed through histology (Figure 5C,D). We observed that the diameters from the simulation scaled linearly to match the diameters observed from histology, with a scaling factor of 0.726 yielding an r-squared value of 0.82 (Figure 5E). The depth did not scale linearly well, however, as the best linear scaling of depth yielded an r-squared value of 0.13 (Supplementary Figure 5). From examining the residuals, we observed that square root transforming the depth before linear scaling would yield a more accurate prediction (Supplementary Figure 5). Indeed, by square root transforming the depth and scaling by a factor of 1.6969 we obtained an r-squared value of 0.41 (Figure 5F). Interestingly, although our model was developed based on histologically derived lesion contours, the resulting model-derived lesion diameters were also highly predictive of OCTA-measured lesion diameters with an r-squared value of 0.60 (Figure 5G). Thus, our quantitative model is able to accurately predict lesion shape, diameter, and depth.

### Large-Scale Neurophysiological Recording Before, During, and After Lesion Formation

A valuable tool for studying neurophysiological dynamics following cortical lesioning *in vivo* is the ability to record neural activity. In monkeys D and E we used a semi-transparent 32-electrode ECoG array to record neural activity in sensorimotor cortex during the formation of a lesion (Figure 6A). The transparency of the ECoG array allowed for the induction of a lesion by illuminating through the array. For monkeys D and E, only a single region in the center of the cranial window was illuminated with a 1.5 mm diameter aperture to induce photothrombosis unilaterally. Similarly, the transparency also enabled imaging of the underlying cortical microvasculature by OCTA to confirm the presence of the lesion (Figure 6A,B). We recorded baseline local field potentials (LFPs) for 30 minutes, followed by 30 minutes during the illumination period, and for up to three hours following illumination.

**Figure 6.**
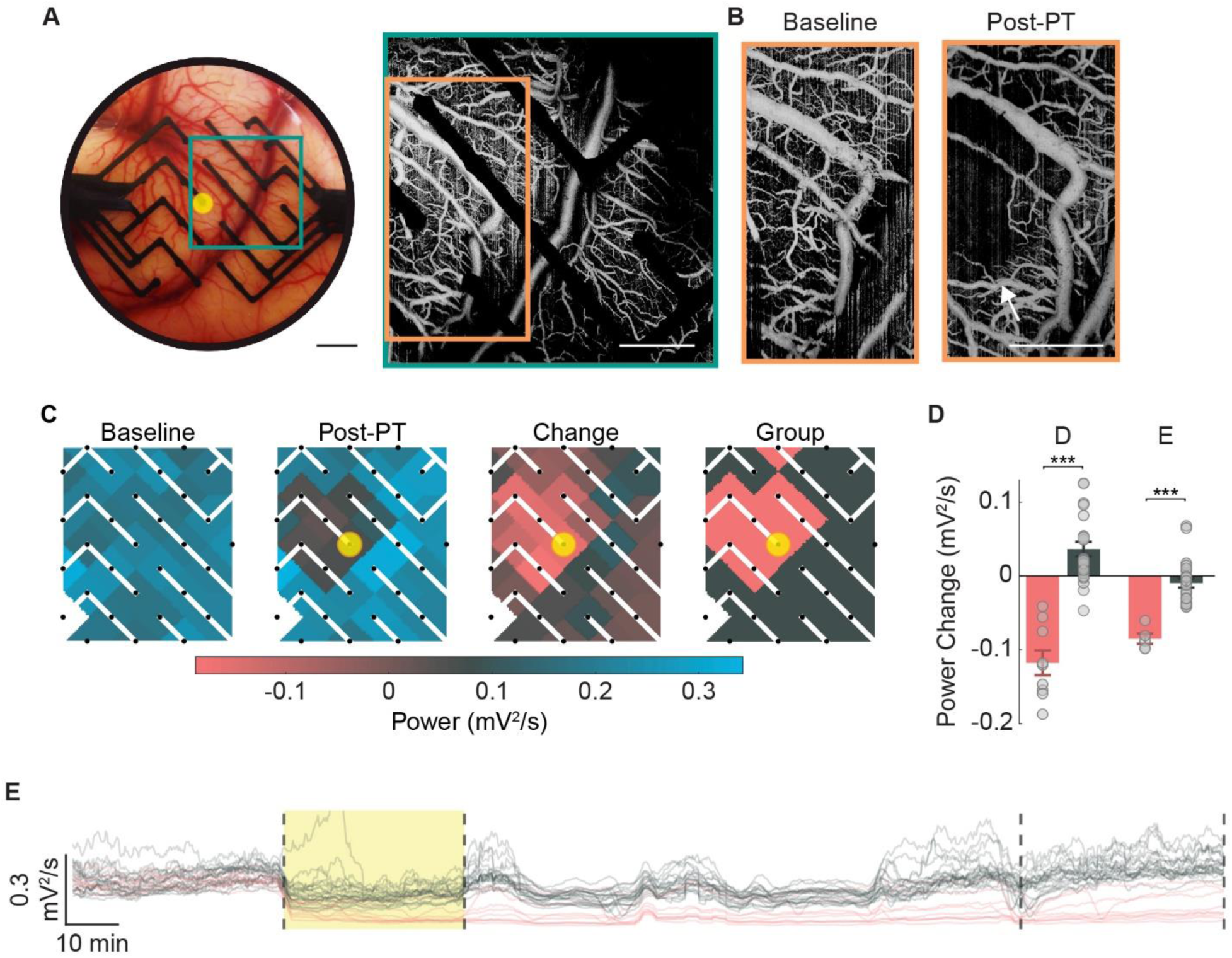
Large-scale electrocorticographic (ECoG) recording of neural activity before, during, and after photothrombotic lesion induction. (A), The transparency of the array enabled OCTA imaging through the array. An example shown here where the area denoted by the green square was imaged in monkey D. Scale bar is 3 mm in the right panel and 2 mm in the left panel. (B), Baseline and 3 hours post-photothrombosis OCTA images demonstrating lesion induction. The imaged areas shown is denoted by the orange rectangle in (A). Scale bar is 2 mm. (C) Gamma band power (30-59 Hz) was calculated from 30 minutes of ECoG recording before and 2.5 hours post-photothrombosis for each channel across the array. The change in power was then calculated for each channel. Channels with statistically significant reductions in gamma power were identified (in pink, paired left-tailed t-test, family-wise error rate ≤ 0.001). Yellow circle indicates the location and extent of the aperture for photothrombotic lesion induction. (D) The average change in power for the lesioned group versus the non-lesioned group (error bars denote ± standard error of the mean, SEM; one-way ANOVA, monkey D: p = 5.2e-9, monkey E: p = 1.0e-5). (E) Example of neural recording (gamma power) through our ECoG array in monkey D as the lesions were induced. Illumination period is shown in yellow. The traces are color-coded according to (D) to show the difference between the lesioned and non-lesioned areas.

To measure the effect of lesioning on neural activity levels, we analyzed the changes in gamma band (30-59 Hz) signal power from LFPs recorded during the formation of the lesions. The average gamma power was calculated over the course of 30 minutes of baseline activity, and between 2.5 to 3 hours after the illumination period (Figure 6C). We calculated the change in power across the array and identified regions with statistically significant reductions in gamma power as corresponding to the lesioned areas (left-tailed paired t-test, family-wise error rate < 0.001). As expected, channels near the illuminated region exhibited significant decreases in gamma power in both monkeys D and E compared to more distant regions (one-way ANOVA, monkey D: p = 5.2e-9; monkey E: p = 1.0e-5; Figure 6C,D). Notably, we demonstrated the capability to monitor neurophysiological dynamics during lesion formation. Here, the reduction in gamma power was observed early during the illumination period across all channels, after which distant channels returned to baseline levels while channels near the illuminated region remained suppressed (Figure 6E).

### Electrical Stimulation Enables Modulation of Perilesional Neural Activity

The ability to stimulate neural activity in the same paradigm that allows for cortical lesioning is critical for the development of stimulation-based interventions. Similar to monkeys D and E, monkeys F and G underwent unilateral photothrombotic lesioning through a 1.5 mm diameter aperture with ECoG recording of activity. We then stimulated through a single perilesional channel in the ECoG array one hour following illumination for a duration of about one hour, after which we continued recording for approximately one hour. Stimulation occurred in six 10-minute blocks separated by 2-minute recording blocks (Figure 7A). To monitor network dynamics following photothrombotic lesioning and during stimulation, we calculated the change in power with respect to baseline in high gamma (60-150 Hz) and theta (4-7 Hz) bands. We observed an overall decrease in high gamma power across the array throughout stimulation in monkeys F and G. This was different from what we observed in non-stimulated monkeys D and E, in which some channels were increasing in high gamma power at equivalent time points (Figure 7B-D). In both stimulated and unstimulated animals, we observed an overall decrease in theta power across the array (Figure 7B-D). Importantly, we demonstrated the ability to electrically stimulate neural activity in the context of our toolbox along with the ability to monitor subsequent effects on network neural dynamics.

**Figure 7.**
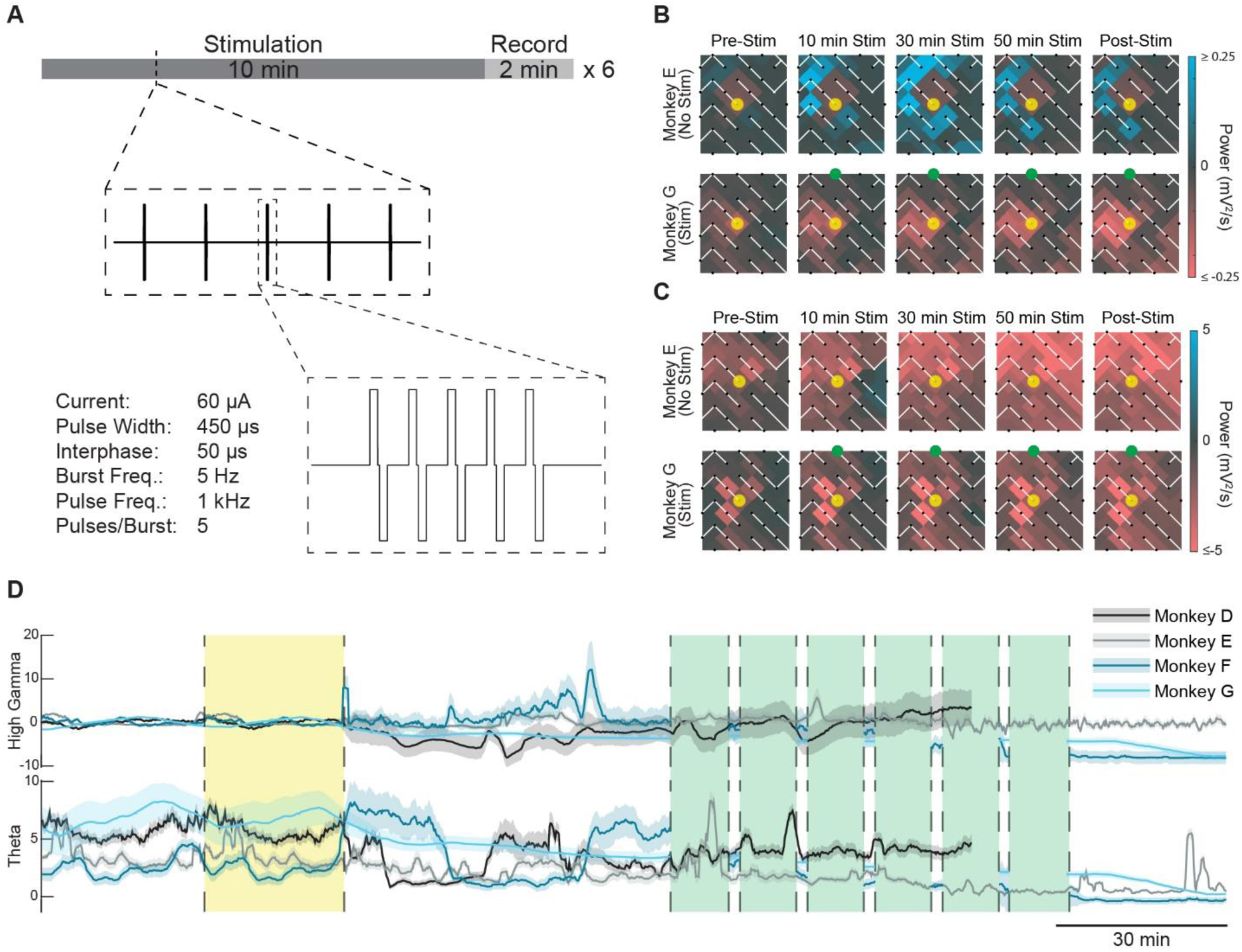
Electrical stimulation enables modulation of peri-lesional neural activity. (A) Stimulation protocol used in this study about one hour after illumination period. Six 10-minute blocks of stimulation were interspaced with 2 min of recording blocks. (B,C) Heat maps of change in high gamma band (B) and theta band (C) power across the ECoG array following blocks of stimulation with respect to baseline for monkeys E (no stimulation) and monkey G (stimulation). Yellow circle indicates the illuminated area and the stimulated channel is indicated by the green node. (D) Average time course of high gamma and theta band power for monkeys D-G. The yellow shaded area indicates the illumination period, while the green shaded areas indicate stimulation blocks for monkeys F and G. Power is normalized to baseline, and the mean power ± 2 x SEM is shown for each animal.

## Discussion

The presented toolbox offers significant advantages for investigating critical questions regarding cortical physiology. Here, we demonstrate the photothrombotic technique for inducing focal ischemic lesions in NHP cortex. The extent of subsequent neuronal cell loss was later confirmed histologically. Moreover, we developed a quantitative model for predicting photothrombotic lesion sizes based on illumination parameters. Through this model, lesions can be designed in accordance with experimental needs. We also present OCTA imaging as a tool for observing vascular dynamics and validating lesion induction *in vivo*. Additionally, our ECoG array allows for the investigation of neural dynamics before, during, and after lesion formation. Critical for developing therapeutic interventions, we also establish the ability to test stimulation-based strategies with our ECoG array. Although not demonstrated here, the toolbox allows for experimentation at clinically relevant time scales when employed in NHPs in contrast to rodent studies.

The photothrombotic technique allows for the induction of targeted focal ischemic lesions in any cortical region of interest. Using the photothrombotic technique, we can achieve greater spatial control over lesion induction in comparison with commonly employed middle cerebral artery occlusion models (Sommer, 2017). The ability to induce consistent smaller, targeted lesions can allow for the induction of lesions to impact specific cortical functions without risking serious injury or mortality compared to middle cerebral artery occlusion (Wu et al., 2016). In rodents, the photothrombotic technique has enabled the study of local cortical remapping in sensorimotor cortex (Harrison et al., 2013). Larger lesions can also be induced with areas as large as the applied cranial window. Moreover, middle cerebral artery occlusion (Maeda et al., 2005), electrocoagulation (Friel et al., 2007; Nudo et al., 1996; Xerri et al., 1998b), and aspiration methods (Padberg et al., 2010) of cortical lesioning require complex surgical intervention and are performed in an invasive manner. In contrast, photothrombotic lesion induction only requires optical access to cortex. With the aid of our predictive computational model, lesions can be designed based on experimental needs. However, as is typical with optical techniques, photothrombotic lesion induction with minimal invasiveness is limited to optically accessible regions. To access subcortical areas, more invasive approaches can be used, such as the use of penetrating optical fibers.

Here, we demonstrated the use of OCTA to image large-scale cortical vascular dynamics and to validate the presence of lesions *in vivo* prior to post-mortem histological analysis. The ability to validate lesion induction *in vivo* is critical for long-term NHP studies. To our knowledge, this is the first time OCTA imaging has been employed to image macaque cortical vasculature to detect the presence of ischemic lesions. We observed strong correlations between OCTA-measured lesion diameters and histologically measured lesion sizes, suggesting that OCTA imaging can be used to reliably measure lesion sizes *in vivo.* Unlike MRI, OCTA imaging enables monitoring of vascular dynamics with greater spatial resolution and without the need for contrast agents, though with limited depth penetration. Moreover, the optical access afforded by our toolbox allows for the use of other optical validation methods, including laser doppler flowmetry (Jiang et al., 2006).

However, not all illuminated regions resulted in lesions detected via OCTA imaging or histology. This may be due to the coincidence of those regions with large blood vessels, in which light penetration through smooth muscle and endothelial cell layers and large lumen diameters prevented the complete occlusion of the vessel. Conversely, successful occlusion of a large vessel may also induce a larger lesion than expected due to the greater extent of downstream or upstream regions. For example, illumination of a larger vessel resulted in the widest lesion observed across our experiments despite the small aperture of 0.5 mm (Figure 4A, monkey C left hemisphere, Supplementary Figure 4). Therefore, illumination of large vessels should be avoided to ensure successful and predictable lesion induction. Moreover, there was one instance in which a lesion was observed via OCTA imaging but was undetected through histology (Figure 4A, monkey B left hemisphere). This may be due to the superficial nature of the lesion such that it was difficult to detect histologically. Consequently, the small size of the lesion may have enabled collateral blood flow to mitigate damage, a phenomenon which has been previously reported in rodent parietal cortex following photothrombotic lesioning of surface vasculature (Schaffer et al., 2006).

Moreover, we demonstrated the ability to monitor underlying neurophysiological dynamics with the ECoG array before and during the development of a lesion for up to three hours. The combination of OCTA imaging and the semi-transparent ECoG array allows for the induction and validation of photothrombotic lesions, the identification of electrodes corresponding with ischemic areas, and the investigation of network physiology and dynamics. Additionally, OCTA imaging and ECoG recording can be used to investigate neurovascular coupling at higher spatial and temporal resolution than functional MRI (fMRI). However, unlike fMRI which allows for online monitoring of neurovascular coupling, OCTA imaging requires *post hoc* offline analysis. The transparency of the array also allows for the incorporation of other optical techniques such as optogenetics and calcium imaging. Simultaneous ECoG recording and optical coherence tomography in combination with optogenetic stimulation has been previously accomplished in mice (Chin-Hao Chen et al., 2020). We demonstrated the ability to test electrical stimulation-based interventions as presented here and in a previous publication (Griggs et al., 2021a), in extension of prior post-lesion stimulation studies (Nudo and Milliken, 1996; Nudo et al., 1996). While we demonstrated the use of this toolbox during acute lesioning, these same tools can be implemented for long-term studies in combination with a chronically implanted cranial window such as our previously published optogenetic interface (Griggs et al., 2019; Khateeb et al., 2019a; WKS et al., 2020; Yazdan-Shahmorad et al., 2015, 2016), which exhibits the same spatial scale. In such studies, new lesions can be created or later enlarged without the need for additional surgical intervention, as would be required with the commonly employed electrocoagulation and cortical aspiration methods of focal cortical lesioning.

We also developed a computational model to predict the shape and scale of photothrombotic lesions. Our regression results and Monte Carlo simulation-based results together allow for planning of precise lesions with a variety of illumination parameters, informed by previous literature and validated against our experimental results. Importantly, through our grid search, we identified the optimal optical properties of cortical tissue to accurately predict lesion sizes. It is important to note, in our experiments we did not vary illumination time across different lesions, with an illumination time based on previous photothrombotic studies (Gulati et al., 2015). Therefore, our computational model was constructed based on lesions induced with consistent illumination times. However, our model may be modified in future studies to predict lesions induced with different illumination times to account for differences in total energy that would be delivered to induce Rose Bengal photoactivation. There are also notable assumptions which are applied to our model. First, we assume a consistent gray matter depth at 2.5 mm, whereas in reality, the depth of gray matter is variable across cerebral cortex, which can affect lesion depth predictions. Moreover, the final step of our model involves scaling the profiles from probability maps to better reconstruct the profiles of our histologically detected lesions. This was done to account for the topography of the cortical vasculature (Figure 5B-D). The depth-wise upscaling of our model contours is likely due to the perpendicular orientation of penetrating arterioles and venules relative to the cortical surface (Gould et al., 2017). The diameter-wise downscaling can be ascribed to the redistribution of blood flow in vessels downstream of an occluded vessel to preserve perfusion in these downstream regions (Schaffer et al., 2006), thus potentially reducing the extent of ischemic damage.

While the presented tools offer great advantages for investigating physiological dynamics in the context of a cortical lesion, there are limitations that can be addressed in later studies. As discussed earlier, the reliance on optical access for lesion induction limits potential regions of interest to superficial brain structures such as cerebral cortex without compromising neural tissue with penetrating optical probes. Additionally, while we developed a model to accurately predict lesion sizes and shapes based on illumination parameters, variations in the induced lesions are likely a result of differences in local optical properties in addition to vascular topographies. Moreover, the ECoG array presented here exhibits limited spatial resolution with 32 opaque electrodes across an area of approximately 314 mm^2^ and an electrode pitch on the order of millimeters. With an array with more electrodes in a more compact arrangement, greater spatial resolution can be achieved. Additionally, while the medium in which in the electrodes are embedded is transparent, the electrodes themselves are opaque, which can occlude optical penetration of cortical tissue for OCTA imaging and photothrombotic lesion induction. Such a concern can be addressed through the use of transparent indium tin oxide electrodes (Ledochowitsch et al., 2015b). While our current ECoG array limits use to LFP recordings, smaller electrodes would allow for the recording of multi-unit activity as has been previously demonstrated (Khodagholy et al., 2015). Furthermore, the potential to induce post-lesion behavioral effects are not addressed in this study. However, the successes of previous macaque studies (Murata et al., 2008; Padberg et al., 2010) with focal lesions of sizes similar to those reported here provide encouraging evidence of the potential to induce behavioral deficits with the photothrombotic technique.

Together, these tools can be used to answer critical questions in neuroscience and drive studies of cortical physiology and related disorders such as stroke and traumatic brain injury. Importantly, our toolbox enables the perturbation of the brain and simultaneous monitoring of physiological dynamics over time. The combination of the presented tools can be used to evaluate vascular dynamics, neural dynamics, and neurovascular coupling in the context of either cortical lesioning, stimulation, or both. Although our toolbox is demonstrated here in an acute setting, the same tools are capable of being used in chronic experiments. Such a capability would allow for the monitoring of the behavioral and physiological effects of lesioning and subsequent recovery over time, in combination with testing stimulation-based interventions. Furthermore, the compatibility of our toolbox with optical tools such as optogenetics can enable the comparison of temporary and permanent cortical lesioning techniques. Importantly, long-term studies employing our toolbox can address key questions regarding cortical functions and drive the development of future rehabilitative therapies for stroke, traumatic brain injury, and other relevant neurological disorders.

## Acknowledgements

We thank Toni Haun, William Ojemann, Evelena Burunova, Stephen Philips, Warren Han, Zhaojie Yao, Teng Liu, Shaozhen Song, Sandi Thelen, Christopher English, Dean Jeffrey, and Britni Curtis for their help with animal surgeries and experimentation. We also thank the Horwitz and Buffalo labs for their shared expertise on histological analysis, and the Ganguly lab for their expertise on the photothrombotic technique. This work was supported by the Eunice Kennedy Shriver National Institute of Child Health & Human Development of the National Institutes of Health (K12HD073945, AY), the National Institute of Neurological Disorders and Stroke of the National Institute of Health (1R01NS116464-01, AY, JZ), the National Eye Institute (R01EY028753, RKW), the Washington National Primate Research Center (P51 OD010425), the University of Washington Royalty Research Fund (AY, KK), the National Science Foundation Graduate Research Fellowship (KK), the Big Data for Genomics and Neuroscience Training Grant (JB), the Weill Neurohub (JZ), and the Center for Neurotechnology (EEC-1028725, DG).

## Author Contributions

KK and JB drafted the manuscript. AY conceptualized the study. AY and KK designed the experiments. AY, KK, DG, MR, JB, JZ, and VK performed the experiments. ML and RKW performed the optical imaging setup and data collection. KK, JZ, MR, JB, and ML performed data analysis. All authors revised and edited the manuscript.

## STAR Methods

### Key Resources Table

| REAGENT or RESOURCE | SOURCE | IDENTIFIER |
| --- | --- | --- |
| **Antibodies** |  |  |
| Mouse monoclonal anti-NeuN | Millipore-Sigma | Cat# MAB377;<br>RRID: AB_2298772 |
| Donkey anti-Mouse IgG (H+L), Alexa Fluor 488 | Invitrogen<br>ReadyProbes | Cat# R37114;<br>RRID:AB_2556542 |
| **Bacterial and virus strains** |  |  |
| **Biological samples** |  |  |
| **Chemicals, peptides, and recombinant proteins** |  |  |
| Silicone base and polymer | Shin-Etsu | Cat# KE-1300T/CAT-1300 |
| Paraformaldehyde, prilled | Sigma Aldrich | Cat# 441244-3KG |
| Cresyl Violet | FD Neurotechnologies, Inc. | Cat# PS102-01 |
| Thionin Acetate salt, Lauth's Violet | Sigma Aldrich | Cat# 861340-5G |
| Triton-X100 | Fisher Scientific | Cat# BP151500 |
| VectaShield mounting medium | Vector Laboratories | H-1000-10 |
| Normal Donkey Serum | Jackson ImmunoResearch Lab | Cat# 017-000-121 |
| Critical commercial assays |  |  |
| Deposited data |  |  |
| Code for power analysis | This paper | N/A |
| Code for photothrombotic quantitative model | This paper | N/A |
| Experimental models: Cell lines |  |  |
| Experimental models: Organisms/strains |  |  |
| *Macaca mulatta/nemestrina* | Washington National Primate Research Center | N/A |
| Oligonucleotides |  |  |
| Recombinant DNA |  |  |
| Software and algorithms |  |  |
| MATLAB R2020a | MathWorks | RRID:SCR_001622,<br>http://www.mathworks.com/products/matlab/ |
| Hugin-2019.2.0 | Hugin Software | http://www.mathworks.com/products/matlab/ |
| Monte Carlo eXtreme (MCX) | Fand and Boas, 2009 | RRID:SCR_007001,<br>http://mcx.sourceforge.net/ |
| Adobe Photoshop | Adobe | RRID:SCR_014199,<br>https://www.adobe.com/products/photoshop.html |
| Other |  |  |

### Resource Availability

#### Lead Contact

Further information and requests for resources and reagents should be directed to and will be fulfilled by the lead contact, Azadeh Yazdan-Shahmorad.

#### Materials Availability

This study did not generate new unique reagents.

#### Data and Code Availability

The data supporting the findings of this study and code used to generate the figures and develop the predictive quantitative model are available from the lead contact upon request.

### Experimental Model and Subject Details

All animal procedures were approved by the University of Washington Institutional Animal Care and Use Committee.

### Method Details

#### Surgical Procedure and Photothrombotic Technique

Three adult *Macaca mulatta* (monkey A, 18.45 kg, 14 years, male; monkey B, 10.75 kg, 16 years, male; monkey C, 10.3 kg, 16 years, female) were anesthetized with isoflurane and placed in a stereotaxic frame. Throughout the procedure we used standard aseptic technique, and the animals’ body temperature, heart rate, electrocardiographic responses, oxygen saturation, and CO_2_ end-tidal pressure were monitored. A sagittal or coronal incision was made to expose underlying soft tissue in the right hemisphere of monkey A and bilaterally in monkeys B and C. After removing the soft tissue to reveal the skull, a 25 mm diameter craniotomy targeting the sensorimotor cortex was made based on the macaque brain atlas (Paxinos et al., 2009) with a center 17.5 mm from the midline and 11.55 mm from the interaural line using a trephine in the right hemisphere of monkey A and bilaterally in monkeys B and C. To allow for optical access to the cortical surface, we performed a durotomy to excise the opaque native dura and replaced it with a transparent silicone artificial dura (10:1 Shin-Etsu KE1300-T and CAT-1300) of 0.5 mm thickness and 25 mm diameter. We acquired baseline OCTA images of the exposed cortical surface through the artificial dura as described later in a later section. Light exposure was limited across the cortical surface by placing on the artificial dura an opaque silicone mask with circular apertures with diameters of 0.5, 1.0, and 2.0 mm distributed across the mask (Figure 1A). We then intravenously injected 20 mg/kg of Rose Bengal solution (40 mg/mL concentration) through the saphenous vein over the course of 5 minutes and began illumination through the mask with a cold white light source for 30 minutes (Figure 1B). To prevent further light exposure following illumination, the cranial window was optically shielded until OCTA images were acquired 3 hours post-illumination. Following post-illumination OCTA imaging, we perfused the animal with 6 L of 4% paraformaldehyde then extracted and immersed the brain in 4% paraformaldehyde solution for 24–48 hours at 4 °C.

### Electrocorticographic Recording, Stimulation, and Power Analysis

#### Electrocorticographic Recording

Four adult *Macaca nemestrina* (monkey D, 12.8 kg, 14 years, female; monkey E, 13.10 kg, 14 years, female; monkey F, 13.8 kg, 14 years, female; monkey G, 14.6 kg, 7 years, male) underwent a surgical procedure similar to that of monkeys A, B, and C. Briefly, bilateral craniotomies and durotomies were performed targeting the sensorimotor cortex following sedation and stereotaxic positioning. After baseline OCTA images were acquired through the transparent silicone artificial dura (monkeys D-E), the artificial dura was removed and replaced with a semi-transparent ECoG array of 32 electrodes (Figure 4A). To correlate electrode location with lesion location, OCTA images were acquired prior to photothrombosis through the array for monkey D (Figure 4A). We placed an opaque mask similar to that of monkeys A through C but with a single aperture located in the center with a diameter of 1.5 mm on the ipsilesional hemisphere (monkeys D, F, and G: left hemisphere; monkey E: right hemisphere). In the contralesional hemisphere we placed a mask without any apertures. A skull screw located anterior and medial to the ipsilesional cranial window was used as a ground for ECoG recordings. Prior to Rose Bengal infusion and illumination as described previously, baseline ECoG recordings were collected bilaterally for 30 minutes. We also recorded LFPs during illumination and for 3 hours following illumination. We then acquired OCTA images in both hemispheres (monkeys D-E), perfused the animals, and stored the extracted brain as described previously.

#### Electrocorticographic Stimulation

One hour following illumination, we stimulated perilesionally through one channel for a duration of about one hour in monkeys F and G. Stimulation (60 µA, 5 biphasic pulses per burst at 1 kHz, 450 µs pulse width, 50 µs interphase interval, 5 Hz burst frequency) occurred in six ten-minute blocks interspaced with 2-minute recording blocks. Post-stimulation recording was then continued for another 30-60 minutes.

#### Power Analysis

LFPs were recorded at a rate of 30 kHz and downsampled to 1 kHz in MATLAB for power analysis. Signals were notch filtered at 60, 120, 180, and 240 Hz, and were bandpass filtered to isolate theta (4-7 Hz), gamma (30-59 Hz), and high gamma (60-150 Hz) bands. After filtering, artifacts were removed from the signal by normalizing the discrete time series of each signal, and samples with an amplitude exceeding 25 standard deviations were identified and excluded from analysis. Similarly, channels with power spectral densities that did not exhibit the expected 1/f^α^ curve were also excluded from analysis. Thus, one channel was excluded from analysis in monkey D, three channels were excluded from analysis for monkey E, and 11 channels were excluded from analysis for monkey F. No channels were excluded for monkey G. The signal power for each frequency band was calculated for each electrode over the course of the baseline and the final 30 minutes of post-illumination recording periods. Signal power was calculated by squaring the filtered signal and dividing by the elapsed time. Change in signal power was calculated by subtracting mean baseline power from the mean post-illumination power. To isolate channels with decreasing power, we performed a left-tailed paired t-test between the 30 minutes of baseline power and the final 30 minutes of post-illumination power calculated at 60 s intervals (family-wise error rate ≤ 0.001 to account for multiple comparisons). Thus, we categorized channels in a lesioned group (decreasing power), and a non-lesioned group (failed to show a decrease in power). A one-way ANOVA was used to compare average change in power of channels across the two groups at a significance level of 0.05. The time-varying signal power across all recording periods was calculated in 10 s intervals and smoothed (smoothing factor of 0.4-0.5). To observe the effects of electrical stimulation over time, the power time-course for each individual channel was normalized to baseline mean and standard deviation.

### Histology and Immunohistochemistry

After 24-48 hours of post-fixation in 4% paraformaldehyde brains of all animals were dissected into a single 25 mm thick coronal block containing the region of interest using a custom matrix and stored in 30% sucrose in PBS solution at 4 °C for a minimum of one week. The block was then frozen and sectioned into 50 µm coronal sections using a cryostat (Microm, Thermo Fisher) or a sliding microtome (Leica). Cut sections were stored in PBS solution with 0.02% sodium azide at 4 °C. To evaluate the extent of ischemic damage to the neuronal population we first performed Nissl staining on mounted serial coronal sections with a rostrocaudal separation of approximately 0.45 mm using Cresyl Violet (monkeys A and B) or Thionin acetate (monkeys C, D, and E). Due to inadequate storage conditions resulting in loss of tissue, not all putative lesions were confirmed histologically for monkey A. Therefore, data from monkey A was excluded from the rest of our analysis. To additionally confirm the neuronal cell loss in the illuminated regions, we immunostained adjacent sections for a selective neuronal marker NeuN. Sections were further trimmed to include the mediodorsal part of the hemisphere with putative lesion sites; these were first incubated in 50% alcohol for 20 min to permeabilize, then rinsed in PBS and incubated in 10% normal donkey serum in PBS containing 0.1% triton-X100 at 4 °C for 1-2 hours, and then incubated with a mouse anti-NeuN primary antibody (1:500, Millipore Sigma; MAB377, RRID: AB_2298772) at 4 °C for 48 hours. Sections were then rinsed in PBS, incubated in 2% normal donkey serum followed by the secondary donkey anti-mouse antibody conjugated to AlexaFluor 488 (1:300, Millipore) for 4-6 hours at room temperature. Sections were rinsed and mounted using VectaShield mounting media (Vector) containing DAPI and imaged using Nikon 6D widefield automated microscope system (Nikon imaging Center at UCSF).

### Lesion Reconstruction and Size Measurement

All Nissl-stained slices for monkeys B through E were imaged and co-registered in MATLAB (MATLAB R2020a, MathWorks) without resizing. The registered images were then edited in Adobe Photoshop to enhance boundaries between infarct and non-infarcted regions, after which each slice was smoothed with an edge-preserving filter, binarized, and underwent edge-detection to identify slice boundaries with and without the infarcted regions in MATLAB (Figure 2D,E). Lesion boundaries were obtained by subtracting boundaries with the lesions from boundaries that excluded the lesions. These boundaries were then visualized as surfaces in three-dimensional space with linear interpolation of the approximately 0.45 mm rostrocaudal separation between slices. The average diameter and maximum depth from representative coronal slices of each lesion were then calculated based on image resolution.

### Optical Coherence Tomography Angiography Imaging

For *in vivo* validation of disrupted blood flow in cortical vasculature, we acquired OCTA images for monkeys B through E. OCTA was acquired from a custom-built prototype OCT system, using a 200 kHz vertical cavity surface-emitting swept source (SL1310V1-10048, Thorlabs Inc., Newton, NJ) with a central wavelength of 1310 nm, and a sweeping bandwidth of 100 nm. The prototype was similar to the device reported in a previous publication (Deegan et al., 2018b; Xu et al., 2017).

OCTA was performed in repeated raster-scan. The scanning beam was directed into the cortical tissue through a transparent silicone artificial dura, prior to photothrombosis and 3 hours post-illumination with a lateral FOV of approximately 9 mm x 9 mm. At each position on the raster scan, one full A-scan approximately 8 mm deep (1408-sampling-pixel) was acquired. Each B-frame contained 1000 A-line scans and was repeated 8 times, then moved to the next B-frame. This process was repeated to form a volume of 1408 x 1000 A-lines x 8 repeat x 1000 B-frames. The optimal resolution in the axial direction (into the tissue) was 5.5 µm and the lateral resolution was 24 µm. The volume scan was repeated at different regions within the cranial window, then processed to form a 3D OCTA data set. For visualization, the 3D OCTA was color-coded with the depth associated with each vessel into a 2D-en-face (top-down) projection.

The 2D-en-face projections at different regions were then stitched together to form a larger FOV as in Figure 3 using the Hugin software (Hugin-2019.2.0). Stitched OCTA images were then aligned with images of the cortical surface via control point registration in MATLAB. To estimate the diameters of the lesions, images were first smoothed with a Gaussian filter and binarized. Lesion boundaries of each detected lesion were fitted with ellipses and the major and minor axis lengths were recorded. The major axis lengths were then defined as OCTA-measured lesion diameters. Illuminated regions outside the FOV excluded from analysis.

### Light Intensity Measurements

Due to the uncollimated nature of the illuminating light beam, the power density of the light through each aperture was assumed to differ based on the location and diameter of each aperture. We measured the light intensity (535 nm) through each aperture (10-20 times) and through the mask without any apertures using a photodetector (Thorlabs PM100D Power Meter and S121C Sensor).

### Lesion Size and Illumination Parameter Analysis

The relationship between OCTA-measured lesion diameter and histologically measured lesion diameter was assessed by fitting an ordinary least squares linear regression in MATLAB (fitlm function). Similarly, the relationships between OCTA-measured diameter and histological lesion depth, as well as histological depth and diameter were analyzed by ordinary least squares regression. We also modeled the combined effects of both aperture diameter and the log-transformed light intensity as predictor variables of each lesion size metric (OCTA-measured diameter, histological diameter, and histological depth) separately in MATLAB by fitting a multivariate ordinary least squares linear regression (fitlm function). To better visualize the multivariate regressions, we plotted partial regressions, in which the adjusted variables represent the mean of the plotted variable plus the residual of the variable fit with the excluded predictor variable (Figure 4G-L).

### Light Simulation

Our computational model of light penetration and subsequent lesion in the brain consisted of a Monte Carlo simulation of photons propagating through brain tissue, extraction of a light intensity contour, and scaling of the contour. A virtual volume was constructed which mimicked the experimental setup and had the relevant optical properties of brain tissue. Photons were virtually propagated through this volume to obtain a probability distribution of photon fluence, or light energy passing through a given area. Optical parameters were informed by published literature regarding primate cortical optical properties and then refined to maximally match our experimental results obtained through histology. The Monte Carlo simulations were designed using the Monte Carlo eXtreme (MCX) software (Fang and Boas, 2009) and run on discrete graphics cards (Nvidia GTX 1080Ti and Nvidia GTX 2060).

The virtual volume consisted of a light source and mask, an artificial dura, and gray and white matter (Figure 5A). The entire volume was 8 mm x 8 mm x 5.6 mm (width x width x height), with voxel resolution of 0.01 mm^3^. We specified the light source as a disk parallel to the surface of the brain with collimated light exiting the side of the disk facing the brain, and then passing through a thin highly scattering disk which uncollimated the light. We set the diameters of both disks to be equal to the aperture diameter of the mask of the corresponding *in vivo* experiment. We specified the volume peripheral to the disks as a highly absorbing and highly scattering medium to mimic the opacity of the non-aperture part of the mask (Supplementary Table 2). The diameter used in our simulations were 0.5 mm, 1 mm, 1.5 mm, and 2 mm, corresponding with the apertures tested in our *in vivo* experiments. After specifying the volumes, we ran simulations of 1 million photons through the volume. The simulations yielded voxel-wise fluence values, or values of radiant energy received per unit area. The fluence results were imported into MATLAB and analyzed with custom code.

Research on the optical properties of gray matter is often contradictory, and values of index of refraction, anisotropy coefficient, and absorption and scattering coefficients vary by orders of magnitude in published research. We defined a grid search boundary of optical properties by using the values reported in previous literature (Gottschalk, 1992; Yaroslavsky et al., 2002) as upper and lower bounds of the properties. As the absorption spectrum for Rose Bengal peaks at 559 nm, we used the optical properties reported for this wavelength by interpolating from nearest reported wavelengths. The range of the grid was [0.03, 0.76] mm^-1^ for gray matter absorption, [9.9, 53.6] mm^-1^ for gray matter scattering, [0.09, 0.36] mm^-1^ for white matter absorption, and [41.9, 78.4] mm^-1^ for white matter scattering. For anisotropy coefficients the two publications reported similar values, therefore we averaged the values between the publications to obtain an anisotropy coefficient of 0.92 for gray matter and 0.8 for white matter. The refractive index was set as 1.36 for gray matter and 1.38 for white matter^54^. From the optical property ranges we constructed a 5 x 5 x 4 x 4 size grid with 5 values for the gray matter absorption and scattering coefficients and 4 values for the white matter absorption and scattering coefficients. For each of the 400 combinations of parameters, four simulations were run, corresponding with the four aperture sizes tested in our experiments. Each simulation yielded a volumetric fluence distribution. We further obtained representative 2D central slices to compare with histologically obtained lesions.

For each photon intensity distribution corresponding to an optical property combination, we identified the light intensity threshold which yielded contours most closely matching those obtained from central slices of lesions obtained from histology. By treating the maximum depth and average width of an individual contour as the prediction of maximum depth and average width of the corresponding lesion, we quantified the predictive power of the simulation. Denote by *SSR_d_* the residual sum of squares of prediction of maximum depth of a lesion, *SST_d_* the total sum of squares of maximum depth of a lesion, *SSR_w_* the residual sum of squares of prediction of average width of a lesion, and *SST_w_* the total sum of squares of average width of a lesion. The objective function used to find the best light intensity threshold was:

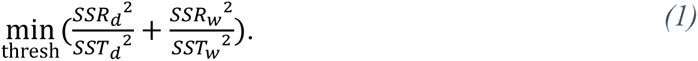

This is equivalent to maximizing the (squared or unsquared) Euclidean norm of the r-squared values of depth and width predictions less 1. Denote by 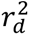 the r-squared value of the simulation-based prediction of maximum depth of a lesion, and 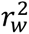 the r-squared value of the simulation-based prediction of average width of a lesion. Then the equation above can be rewritten as:

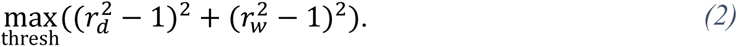

We solved this objective function individually for each optical property combination using Bayesian optimization. We then identified the simulation yielding the minimum value of this objective function as that most closely matching our histology data and used the results of that simulation for all later parts of the analysis. The optical properties of this best simulation were gray matter absorption coefficient of 0.395 mm^-1^, gray matter scattering coefficient of 53.6 mm^-1^, white matter absorption coefficient of 0.09 mm^-1^, white matter scattering coefficient of 54.066 mm^-1^, and a light intensity threshold for lesion induction of 19.9 µW/mm^2^.

We then identified a transformation to accurately scale the profiles of the light simulation contours to that of the lesions obtained from histology (Figure 5C,D). We opted to do this by independently scaling the width and depth of the contours. Our first approach was to fit a univariate linear regression from the simulation depths and widths to the lesion depths and widths, respectively. In this linear regression we did not allow an intercept term. Based on the results of our linear regressions, we further evaluated the need for non-linear transformations. Once these scaling factors were set, we treated them as the final stage of our modeling platform, whereby light intensity contours from the Monte Carlo would be stretched and shrunk accordingly to transform from the light profile to the lesion profile.

## Supplemental Information Titles and Legends

**Supplementary Figure 1.**
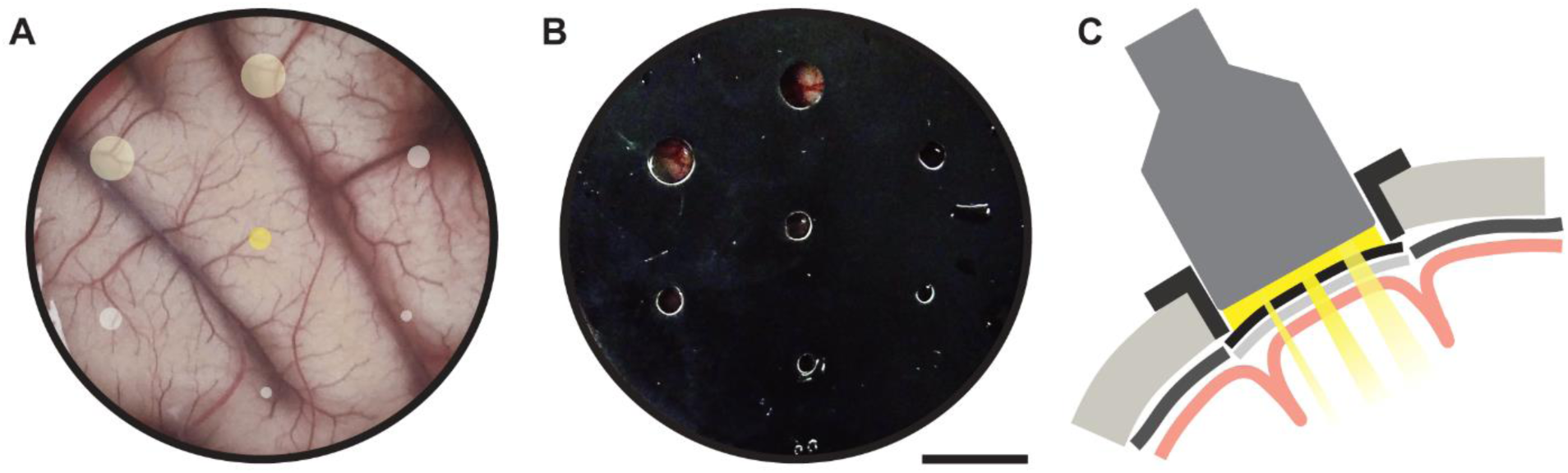
Schematic of illumination using a multi-apertured mask used for monkeys A, B, and C. (A) A cranial window through an artificial dura with projected areas of illumination shown in yellow. (B) Multi-apertured mask is placed on top of the artificial dura. (C) Coronal schematic of illumination through the apertured mask. This setup enables testing different illumination intensities and aperture diameters.

**Supplementary Figure 2.**
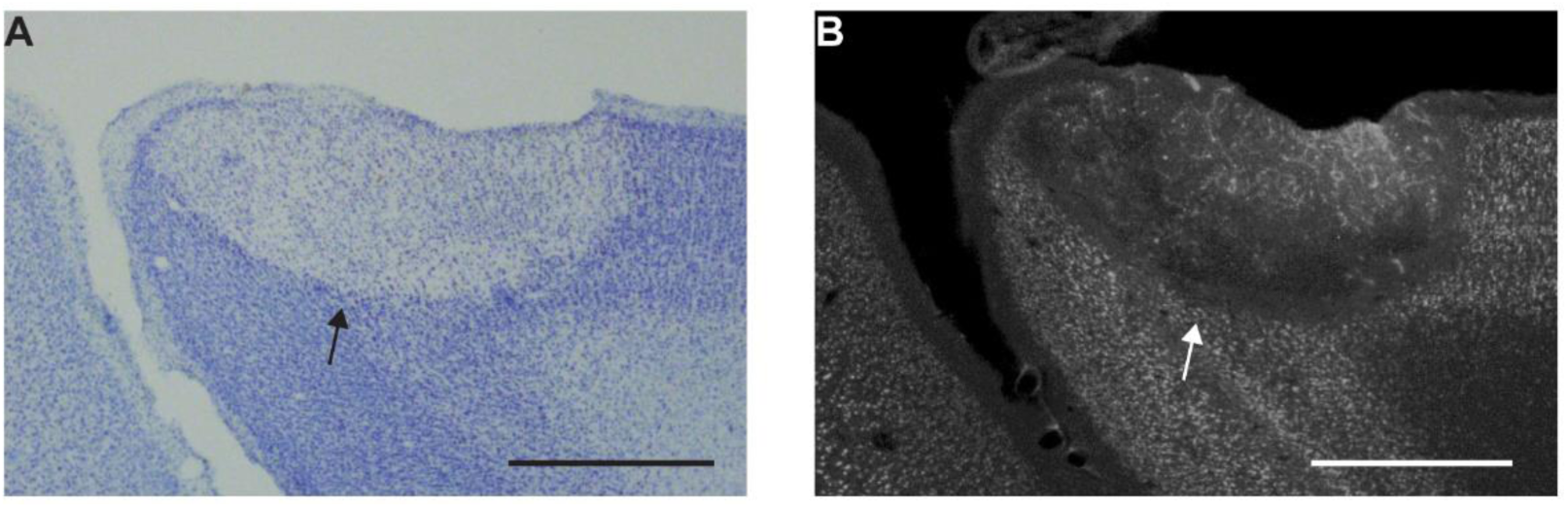
Validation of neuronal cell loss with NeuN staining. (A) Nissl staining demonstrating cell loss indicated by the black arrow in a representative coronal slice from monkey C. (B) NeuN staining of an adjacent slice demonstrating neuronal cell loss in the same lesion indicated by the white arrow. Scale bars are 1 mm.

**Supplementary Figure 3.**
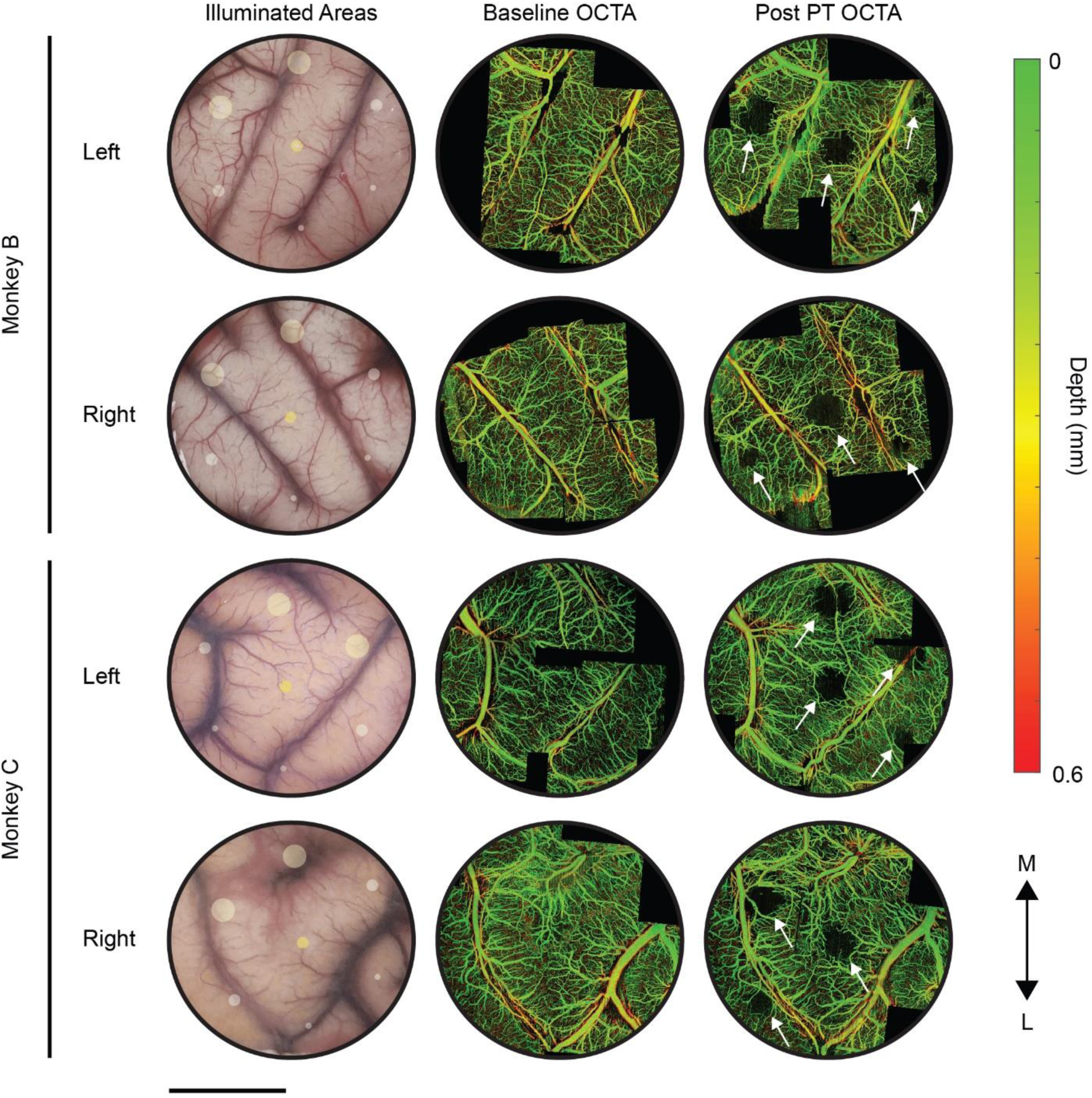
Optical coherence tomography angiography (OCTA) imaging was used to validate lesion induction *in vivo*. Stitched baseline and post-photothrombosis OCTA images are shown for both hemispheres of monkeys B and C, with vasculature color indicating cortical depth. In the left column, yellow circles indicate the illuminated regions for photothrombosis, with the diameters corresponding with the aperture diameters. Shading of the yellow circles in the left column indicate relative intensity of illumination. Baseline OCTA images are shown in the middle column. In the right column, dark regions indicated by white arrows in OCTA images taken 3 hours after photothrombosis indicate successful lesion induction due to the occlusion of the illuminated microvasculature. Scale bar is 1 cm.

**Supplementary Figure 4.**
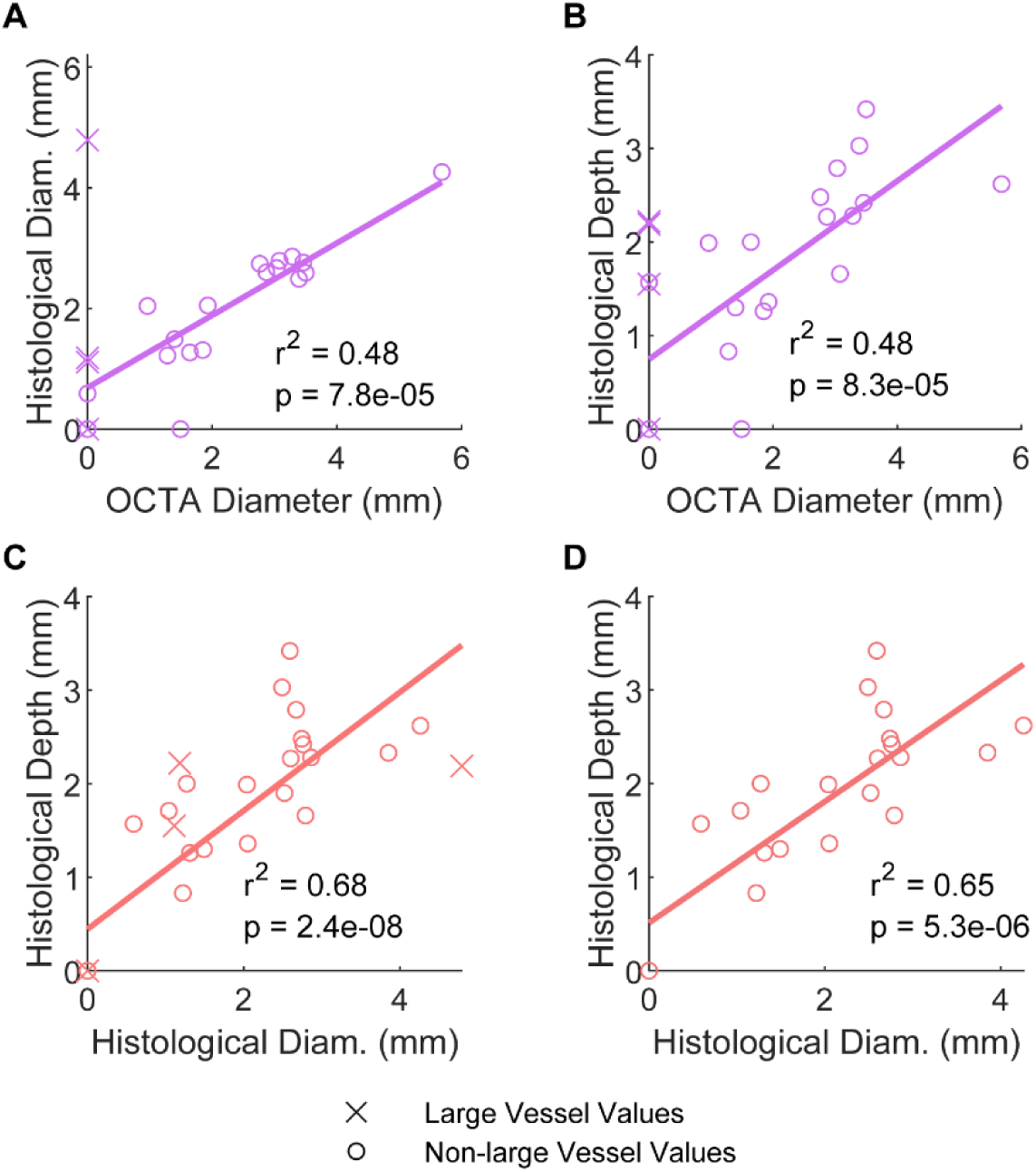
OCTA predictions of histologically measured lesion size with large vessel values included. (A) Correlation between OCTA- and histologically measured lesion diameters (0.49 r-squared, p = 7.8e-5). (B) Correlation between OCTA-measured lesion diameters and histologically measured lesion depths (0.48 r-squared, p = 8.3e-5). (C,D) Correlation between histologically measured lesion diameters and depths with large vessel values included (C, 0.68 r-squared, p = 2.4e-8), and excluded (D, 0.65 r-squared, p = 5.3e-6).

**Supplementary Figure 5.**
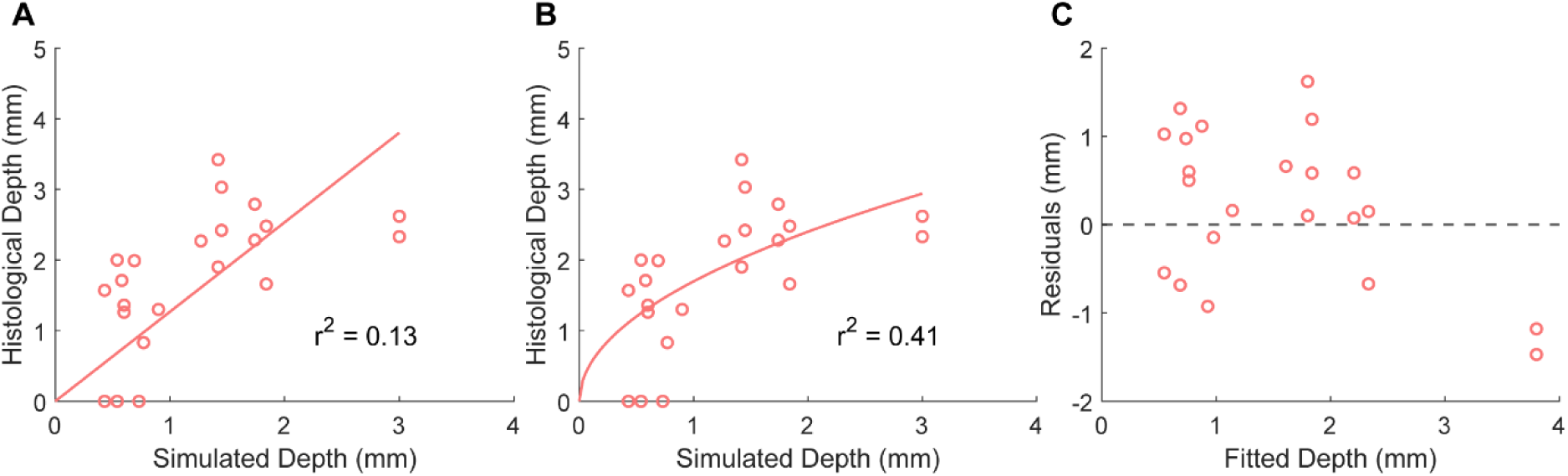
Transformation of simulated depths in second stage of modeling photothrombotic lesions. (A) Histological depths vs linearly transformed simulated depths. (B) Histological depths vs square root square root transformed simulated depths. (C) Residuals of histological depths vs linearly transformed simulated depths plotted against fitted values from (A).

**Supplementary Table 1.** Table of the estimated coefficients and corresponding p-values for multivariate prediction of each lesion size metric.

|  | OCTA Diameter |  | Histological Diameter |  | Histological Depth |  |
| --- | --- | --- | --- | --- | --- | --- |
|  | Estimate | p Value | Estimate | p Value | Estimate | p Value |
| Intercept (mm) | 2.0381 | 0.00016829 | 1.9002 | 3.5227e-05 | 1.2064 | 0.016778 |
| Log(Intensity (mW)) | 0.57968 | 7.7597e-06 | 0.46653 | 2.23e-06 | 0.20399 | 0.036939 |
| Aperture Diameter (mm) | 0.90238 | 0.0066161 | 0.55467 | 0.036591 | 0.71685 | 0.037535 |

**Supplementary Table 2.**
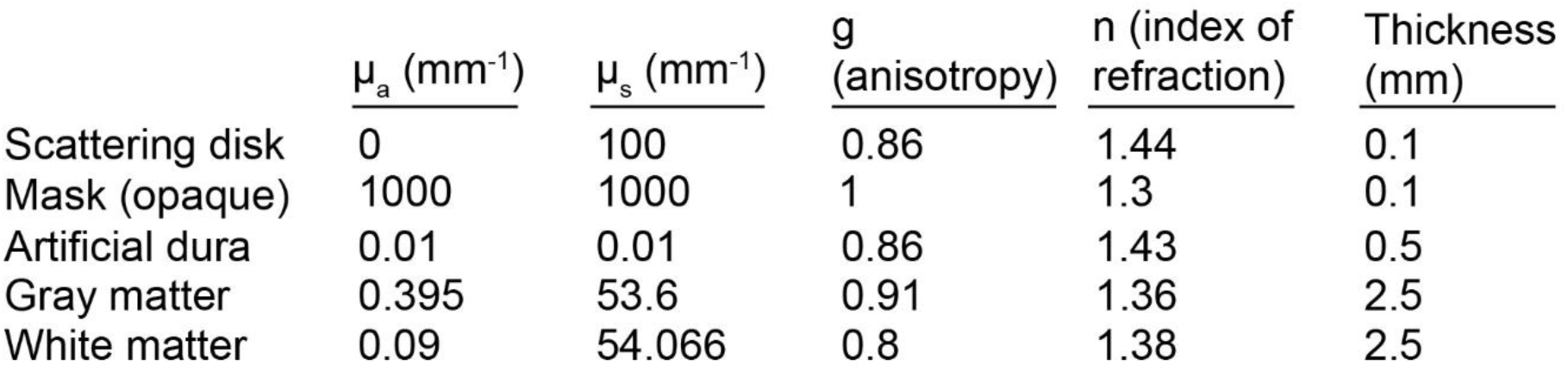
Table of optical properties and thicknesses of virtual media used in the Monte Carlo simulation.

| | $\mu_a$ (mm<sup>-1</sup>) | $\mu_s$ (mm<sup>-1</sup>) | g (anisotropy) | n (index of refraction) | Thickness (mm) |
| --- | --- | --- | --- | --- | --- |
| Scattering disk | 0 | 100 | 0.86 | 1.44 | 0.1 |
| Mask (opaque) | 1000 | 1000 | 1 | 1.3 | 0.1 |
| Artificial dura | 0.01 | 0.01 | 0.86 | 1.43 | 0.5 |
| Gray matter | 0.395 | 53.6 | 0.91 | 1.36 | 2.5 |
| White matter | 0.09 | 54.066 | 0.8 | 1.38 | 2.5 |

